# Infectious bursal disease virus suppresses H_9_N_2_ avian influenza viral shedding in broiler chickens

**DOI:** 10.1101/378703

**Authors:** Vahid Reza Ranjbar, Ali Mohammadi, Habibollah Dadras, Arash Bidadkosh

**Author notes:** These authors contributed equally to this work (Ali Mohammadi, Habibollah Dadras).

## Abstract

Infectious bursal disease (Gumboro) virus causes immunosuppression in chickens, increasing their susceptibility to other viral and bacterial diseases and resulting in vaccination failure. In the present study, we investigated the immune-depressing effect of infectious bursal disease virus on H9N2 avian influenza viral infection in commercial broiler chickens. Chickens were divided into four groups. In group A, chickens were inoculated with Gumboro virus at 21 days of age and H9N2 influenza virus 5 days later. Groups B and C only received influenza virus at 26 days of age and Gumboro virus at 21 days, respectively. Chickens in the control group (D) were inoculated with normal saline at the same times.

Tissue samples from different organs were collected on days 1, 3, 6, 9, and 12 after H9N2 infection. Macroscopic observation showed bursal disease lesions in groups A and C, including swollen bursa with the presence of gelatinous exudates, hemorrhages in the thigh muscle, edema, and nephritis.

Reverse transcription polymerase chain reaction was used to study H9N2 influenza virus dissemination, and quantitative reverse transcription polymerase chain reaction to determine viral genome copy number in different organs. A considerable titer of avian influenza virus was found in the trachea, lungs, cecal tonsils, spleens, and feces of infected chickens. The genome copy number of the virus in the trachea and lungs of group A was significantly higher than that in group B on the first day after inoculation. However, our method did not detect the avian influenza virus genome in group A. In conclusion, we suggest that pre-exposure to Gumboro virus at 3 weeks of age reduces the replication and shedding of H_9_N_2_ in broiler chicken.

## Introduction

Infectious bursal disease virus (IBDV) is a member of the Birnaviridae family, which has double-stranded RNA genomes. The virus results in acute, highly contagious infections, immunosuppression, and mortality in young chickens [1, 2], and related economic losses to chicken farms are substantial. Bursal damage is the main feature of the disease (IBD or Gumboro) and is observed in affected chickens worldwide [1]. In IBD, immunosuppression increases susceptibility to other viral diseases, such as viral arthritis [3], Marek’s disease [1], infectious bronchitis [4], infectious laryngotracheitis [5], inclusion body hepatitis [6], and chicken infectious anemia [7], as well as infection by protozoa [8].

The H_9_N_2_ avian influenza virus (AIV) is a pathotype of low pathogenic avian influenza (LPAI) virus in broiler chickens [9] and has been reported in Asia, especially in the middle east [10, 11]. Subtypes of this virus are now known to be enzootic in rural areas of Iran and present in commercial chicken farms. Although H_9_N_2_ AIV is an LPAI virus, it has severe pathogenicity and causes a high mortality rate among farm chickens co-infected with other bacterial and viral diseases [9, 12]. As IBDV is involved in many such co-infections, a greater degree of influenza viral replication is expected in IBDV-infected chickens. To determine to what extent IBDV affects sequela of H9N2 infection, we studied the effects of IBDV inoculation on subsequent H9N2 AIV infection in 215 broiler chicken, in term of clinical signs, gross pathology, serology, virus dissemination in different organs, and shedding of the virus from trachea as well as in the feces.

## Materials and methods

### Birds

A total of 215 one-day-old Cobb broiler chickens were purchased from a local hatchery and reared in the experimental research unit of the Shiraz University College of Veterinary Medicine (Shiraz, Iran). All birds were kept under controlled conditions based on the Cobb breeder management catalog and had free access to food and water throughout the experiments. The chickens received no vaccinations. The study protocol, including animal sacrifice, was approved by the research ethics committee of Shiraz University. In addition, we followed and adapted the recommendations of the European Council Directive (86/609/EC) of November 24, 1986, regarding standards for the protection of animals used for experimental purposes. To determine maternal immunity ten 3-day-old chickens were randomly selected and euthanized. Blood samples were then collected and tested for serum antibodies against IBDV and H9N2 AIV using ELISA and hemagglutination inhibition (HI), respectively.

### Experimental design

At 21 days of age, the chickens were inoculated with IBDV. One day before inoculation, five chickens were randomly selected and blood samples taken from their wing veins. The birds were then euthanized by cervical dislocation. On necropsy, tissue samples of the trachea and the bursa of Fabricius were immediately collected and preserved at -80°C until analyzed. Samples were subsequently tested for possible contamination with Newcastle disease virus, AIV [13], IBDV [14], and infectious bronchitis virus using RT-PCR analysis [15]. Negative results were required before starting the inoculations. The other birds, including two hundred broiler chickens, were randomly selected and divided into two equal groups (I and II) of 100 chickens each and were reared in a separate, isolated room under controlled conditions. For induction of IBD, we used isolates of Shiraz IBDV (Accession number: JX983160), a highly virulent infectious bursal disease virus enzootic in the region. After a 3 h fast, chickens in the treatment group (group I) were infected with 10^4.2^ ELD_50_/0.5 ml of vvIBDV isolate via intra-crop inoculation. The control group (group II) chickens only received 0.5 ml of normal saline. On day 3 after inoculation, 5 chickens from both groups were euthanized, and blood samples were collected. Tissue samples from the trachea and the bursa of Fabricius were then isolated and frozen in liquid nitrogen at - 80°C. The tissue samples were prepared and tested by RT-PCR analysis to determine the presence or absence of IBDV infection in the chickens. Trachea tissue and feces were analyzed by RT-PCR to look for AIV contamination. In addition, serum samples were collected from both groups and tested by HI to confirm the RT-PCR results of AIV contamination.

On day 5 after inoculation, chickens in the treatment group were assigned to subgroup A (70 chickens) or C (25 chickens), and chickens in the control group were divided into subgroups B (70 chicken) and D (25 chickens). In the AIV inoculation step, each group was transferred into a separate, isolated room. Chicken in subgroups A and B were then intranasally inoculated with 10^65^ EID_50_/0.1 ml of A/Chicken/Iran/772/1998(H_9_N_2_) AIV, and subgroups C and D received sterile saline only.

The chickens were observed for any clinical signs of avian influenza (AI) twice daily. In each subgroup, five chickens were randomly selected and euthanized 1, 3, 6, 9, and 12 days post-inoculation (dpi). Tissue samples from the trachea, lungs, brains, spleens, cecal tonsils, and feces were then isolated and preserved at -80°C until analyzed. In addition, blood samples were collected from each subgroup and tested for antibodies against influenza virus by an HI assay. To trace the possible H9N2 viremia in chickens from subgroups A and B, samples were collected from 5 euthanized chickens every 8 h for 3 dpi. Buffy coats were isolated from the blood samples and tested for the presence and quantity of virus using RT-PCR.

### RNA extraction

Total RNA in the buffy coat samples was isolated using a Cinnapure RNA kit (CinnaGen Molecular Biology and Diagnostic, Tehran, Iran) according to the manufacturer’s instructions. Briefly, 400 μl of lysis buffer was added to 100 μl of the sample, and the solution was mixed. Then, 300 μl of precipitant solution was added, and the solution was mixed again. A total volume of 800 μl of solution was transferred into the included column and centrifuged at 12000 rpm for 15 min. The columns were washed twice with the washing buffer. Total RNA was dissolved in 30 μl of RNase-free water via centrifugation and stored at -70°C.

Tissue and fecal samples were prepared for RNA extraction using RNX-plus solution (CinnaGen). Briefly, a 10% (W/V) suspension of feces was prepared in DEPC water and centrifuged at 250 *g* for 10 min at 4°C. The supernatant was removed, and tissues weighing 50–100 mg were then homogenized in RNX-plus. RNA was isolated according to the manufacturer’s protocol. All RNA samples were normalized to the lowest concentration in DNase-/RNase-free distilled water before quantitation.

### Purification of recombinant plasmid

*Escherichia coli* XL1-blue containing a plasmid ligated with a partial sequence of the avian influenza M gene was cultured in 5 ml of LB medium for 24 h. The medium was then centrifuged at 250 *g* for 10 min, and precipitated bacteria were subjected to plasmid isolation using an AccuPrep Plasmid Extraction kit (Bioneer, Daejeon, Korea) according to the manufacturer’s instructions. The number of isolated plasmids was measured in solution using a spectrophotometer (BioPhotometer; Eppendorf, Hamburg, Germany). The recombinant plasmids were used as standards in the quantitation assay.

### Reverse transcription PCR

For the synthesis of cDNA, 5 μl of isolated RNA, 1 μl containing 20 pmol of H_9_ forward primer (AccuPower RT PreMix kit; Bioneer), and 14 μl of DEPC distilled water were added and mixed in a lyophilized tube. cDNA synthesis was carried out at 45°C for 1 h and 95°C for 5 min. Isolated cDNA was then collected in tubes and stored at -20°C until analyzed.

The total reaction volume was 20 μl, and comprised 2 μl of 10x PCR buffer, 0.6 μl of MgCl_2_ (1.5 mM), 0.4 μl of dNTPs (0.2 mM), 0.2 μl of Taq DNA polymerase (1 U), 1 μl of each primer (10 pmol), 9.8 μl of distilled water, and 5 μl of cDNA. The PCR program used with the MJ thermal cycler (Bio-Rad, Hercules, CA, USA) began at 94°C for 5 min, and included 35 cycles at 95°C for 1 min, 53°C for 1 min, and 72°C for 1 min. Final elongation of the cDNA was performed at 72°C for 10 min [16]. PCR products (488 bp) were then visualized in 1% agarose gel, following 45 min electrophoresis at 100 V.

### Real-time PCR

The total volume of the reaction was 20 μl, and it comprised 2 μl of 10x PCR buffer, 0.4 μl of dNTPs (0.2 mM), 2.4 μl of MgCl2 (6 mM), 0.2 μl of Taq DNA polymerase (1 U), 1 μl of each primer (10 pmol), 0.6 μl of TaqMan probe (6 pmol), 7.4 μl of distilled water, and 5 μl of cDNA. The PCR program used with the Bio-rad Miniopticon system was run at 95°C for 5 min, and included 42 cycles at 95°C for 15 sec and 60°C for 1 min [11]. Specific sequences of the isolated oligonucleotides are shown in Table 1.

**Table 1.**
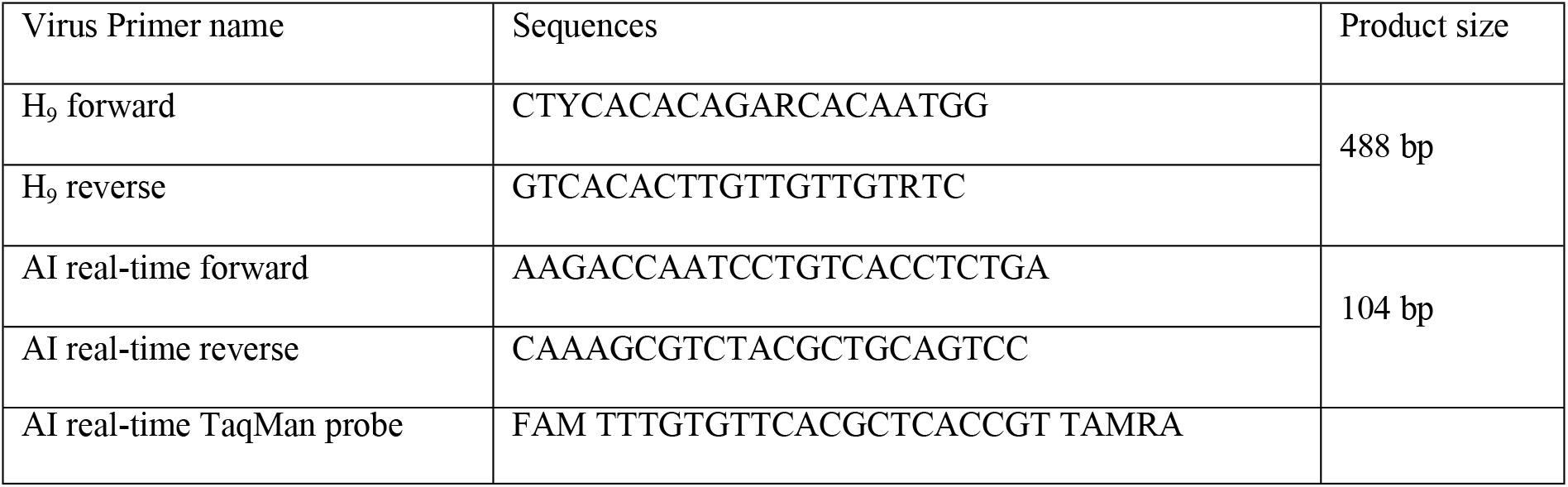
The primers and TaqMan probe sequences used in reverse transcription polymerase chain reaction and real-time polymerase chain reaction to detect and measure genome copy number of H_9_N_2_ avian influenza.

### HI test

To evaluate the serological status of chickens in different groups against the H9 antigen, sera samples were collected from chickens aged 3, 20 (the day before IBDV inoculation), and 25 days. In addition, H_9_ antibody titers against influenza virus were evaluated 1, 3, 6, 9, and 12 dpi. H9N2 AIV antigen equivalent to four hemagglutinating units (HA) was used to determine the HI activity of two-fold serially diluted test sera in a 96-well microtiter plate [17].

### ELISA

The presence of IBDV maternal antibodies was determined in chickens aged 3 days using an IDEXX ELISA kit (Westbrook, ME, USA) according to the manufacturer’s instruction.

### Statistical analysis

Serological analysis results of different groups were compared by two-way ANOVA followed by Tukey’s range test using SPSS version 13 (SPSS, Inc., Chicago, IL, USA; Table 2). A p-value less than 0.05 was considered significant.

**Table 2.** Mean HI antibody titers against H9N2 AIV antigen at time points after inoculation in different groups.

| Subgroups | Mean antibody titers after AIV inoculation |  |  |  |  |
| --- | --- | --- | --- | --- | --- |
|  | Day 1* | Day 3 | Day 6 | Day 9 | Day 12 |
| A | 2.4 ± 0.44<sup>a</sup> | 3.4 ± 0.39<sup>a</sup> | 4.4 ± 0.66<sup>a</sup> | 5.2 ± 0.73<sup>a</sup> | 5.4 ± 0.24<sup>a</sup> |
| B | 2.4 ± 1.29<sup>a</sup> | 3 ± 0.58<sup>a</sup> | 3.4 ± 0.9<sup>a</sup> | 5.6 ± 0.29<sup>a</sup> | 6.2 ± 0.41<sup>b</sup> |
| C | 2.66 ± 0.24<sup>a</sup> | 2.66 ± 0.82<sup>a</sup> | 2.33 ± 1.04<sup>a</sup> | 2 ± 0.44<sup>b</sup> | 2.13 ± 1.36<sup>c</sup> |
| D | 3 ± 1.15<sup>a</sup> | 2 ± 1.2<sup>a</sup> | 2.33 ± 0.24<sup>a</sup> | 2.2 ± 0.29<sup>b</sup> | 2.33 ± 0.94<sup>c</sup> |

\* The mean values in the same columns (vertical) with different superscripts are significantly different ( $p < 0.05$ ).

## Results

### Clinical signs

Before IBDV inoculation, chickens in all groups were observed for clinical signs of any disease or abnormality. Molecular tests were negative for AI, infectious bronchitis, IBD, and Newcastle disease infection. Three days after IBDV inoculation, chickens in subgroups A and C showed clinical signs of depression, ruffled feathers, and white diarrhea. In addition, chickens in subgroup A showed ruffled feathers, depression, cloudy eyes, conjunctivitis, respiratory distress, coughing, sneezing, and gasping 2 dpi, whereas those in subgroup B showed less severe signs of infection at 3 dpi. The most definite signs of infection in the respiratory system were reported at 6 dpi with H_9_N_2_ AIV and observed in chickens from subgroups A and B. However, the clinical signs of infection in chickens from subgroup B disappeared at 9 dpi, while those in subgroup A continued for up to 12 dpi. Chickens in the control subgroup (D) showed no clinical signs of infection during the study.

### Macroscopic Gumboro disease lesions

At 3 dpi, five chickens were randomly selected from subgroups A and C, sacrificed, and macroscopically observed. Gross lesions in these groups included general edematous and swollen bursa, which are associated with the presence of gelatinous exudates, hemorrhages in thigh muscle, and nephritis. In group C, the bursa of Fabricius’ size at 12 dpi was significantly decreased to approximately half the size of an intact bursa with few gelatinous exudates inside. Additionally, mild nephritis was observed in chickens affected by Gumboro disease.

### Macroscopic avian influenza lesions

At 1 dpi with influenza, macroscopic observation of chickens in subgroups A and B showed no lesions, but did reveal some congestion in their spleens. At 3 dpi with influenza, the chickens in these groups showed clinical signs of respiratory infection, including mild airsacculitis, -with the presence of congestion, and transudates in the trachea. The severity of the lesions in these chicks was increased at 6_dpi. Moreover, the bursa was extremely degenerated at this time. Nephritis and splenitis in the chicks of subgroup A were more severe than those of subgroup B. At 9 dpi, congestion, hemorrhaging, and fibrinous casts were observed in lungs isolated from two chickens in subgroup A, whereas pericarditis and perihepatitis were seen in one chicken in subgroup B. Respiratory lesions were decreased in the other birds from these subgroups. There were no necropsy injuries at 12 dpi except a mild tracheal congestion in the chicks of subgroup A. Necropsy revealed no lesions related to AIV infection in chickens from subgroups D (control) or C.

### H_9_N_2_ HI antibody titer

The mean titers of antibodies after H9N2 inoculation are presented in Table 2. The log-2 transformed mean titers before inoculation were 7.2, 4.12, and 3.5 in the blood samples of chickens aged 3, 20, and 25 days, respectively. At 9 dpi, antibody titers had significantly increased in subgroups A and B. In addition, mean titers in subgroup B were significantly higher than those in subgroup A at 12 dpi.

### RT-PCR

Molecular tests were negative for AI, infectious bronchitis, IBD, and Newcastle disease infection one day before IBDV inoculation. To confirm IBDV infection in group I, the bursae of Fabricius were isolated from five chicks in each group and subjected to RT-PCR analyses using VP2 gene primers 3 days after IBDV inoculation. The results showed a 743 bp amplicon, whereas group II, was negative for IBDV infection. All tested samples were negative for AIV infection at this point.

On 1, 3, 6, 9, and 12 dpi with AIV, 5 chickens from each subgroup were euthanized, and their spleens, trachea, cecal tonsils, lungs, brains, and feces were isolated and tested for H9N2 using RT-PCR. Blood samples were isolated at 8 h intervals and tested up to 3 days after influenza infection. The results were negative in subgroups C & D (Table 3). In subgroups A and B, H_9_N_2_ was not detected in buffy coat samples.

**Table 3.** Number of positive samples and mean copy number of H_9_N_2_ avian influenza genome in various tissues of chickens in the experimental groups.

|  | Days after inoculation of H9N2 AI virus |  |  |  |  |
| --- | --- | --- | --- | --- | --- |
|  | 1 | 3 | 6 | 9 | 12 |

| Subgroups<br>Tissue | A | B | A | B | A | B | A | B | A | B |
| --- | --- | --- | --- | --- | --- | --- | --- | --- | --- | --- |
| Spleen | 0/5* | 0/5 | 0/5 | 0/5 | 0/5 | 0/5 | 0/5 | 1/5<br>5.68 x 10<sup>4</sup>** | 0/5 | 0/5 |
| Trachea | 3/5<br>1.2 x 10<sup>5</sup> | 1/5<br>2.64 x 10<sup>4</sup> | 0/5 | 4/5<br>2.4 x 10<sup>5</sup> | 0/5 | 3/5<br>7.5 x 10<sup>4</sup> | 0/5 | 2/5<br>10.4 | 0/5 | 1/5<br>4.4 x 10<sup>2</sup> |
| Cecal tonsil | 0/5 | 0/5 | 0/5 | 1/5<br>1.39 x 10<sup>5</sup> | 0/5 | 3/5<br>5.8 x 10<sup>4</sup> | 0/5 | 4/5<br>1.5 x 10<sup>4</sup> | 0/5 | 2/5<br>3.2 x 10<sup>4</sup> |
| Lung | 1/5<br>3.86 x 10<sup>2</sup> | 0/5 | 0/5 | 0/5 | 0/5 | 2/5<br>2 x 10<sup>4</sup> | 0/5 | 0/5 | 0/5 | 0/5 |
| Feces | 0/5 | 0/5 | 0/5 | 1/5<br>1.32 x 10<sup>5</sup> | 0/5 | 2/5<br>1.4 x 10<sup>4</sup> | 0/5 | 1/5<br>3.64 x 10<sup>3</sup> | 0/5 | 0/5 |
| Brain | 0/5 | 0/5 | 0/5 | 0/5 | 0/5 | 0/5 | 0/5 | 0/5 | 0/5 | 0/5 |

\* = Number of positive samples/total samples taken. \*\* = mean copy number of H<sub>9</sub>N<sub>2</sub> avian
influenza genome.

### Real-time PCR

Samples with positive RT-PCR results were subjected to real-time PCR to quantify their AIV M gene copy numbers. The efficiency of the test in different tissues was approximately 98% using five 10-fold dilutions of the recombinant plasmid. Mean copy numbers of the M segment at different days after inoculation are shown in Table 3.

## Discussion

The strain of virus used for induction of Gumboro disease in the chickens was highly virulent and causes severe lymphoid necrosis and degeneration in the bursa of Fabricius. Sharma et al. (2000) previously reported that immunosuppression by IBDV is achieved when chickens are inoculated at early ages [1]. Since commercial broiler chickens are protected for up to 2–3 weeks by maternal antibodies, immunosuppression by IBDV is a rare occurrence. In the present study, the chickens were inoculated at 21 days. Therefore, we did not expect the immunosuppressive effect of IBDV to be very strong. However, our results showed that HI antibody titer was lower in subgroup A (treated with IBDV) than in subgroup B on day 12 after AIV infection, which may be due to IBDV immunosuppression or lower AIV replication.

In this study, no AIV shedding was observed in subgroup A except at 1 dpi. In subgroup B, in which chickens were inoculated with only AIV, significant increase in viral shedding was observed over the course of the study. These results suggest interference between IBDV and AIV infection. Meng et al. (2011) reported that AI loads were significantly decreased in trachea isolates from specific-pathogen-free chickens fed with IFN-α as compared with those in control chickens. Additionally, in chickens infected by IBDV, cellular immunity is less affected than is humeral immunity [18]. Therefore, we suggested that study of alterations in serum interleukin levels might aid understanding of the mechanism modulating interference between the two viral infections. Raj et al. (2011) reported that serum levels of IL1-β, IL2, IL8, IL12, IL17, and interferon (IFN)-α and -β are significantly increased in commercial chickens infected with IBDV [19].

Knowledge of the quantity and duration of viral shedding from animals can help us clarify ambiguities in pathogenesis as well as the severity and period of the virus spreading in herds, which may be useful in disease control. Infected chickens shed H_9_N_2_ virus via feces more than via respiratory secretions, particularly 5–7 after infection. Although the virus replicates in the kidneys up to 9–11 days after infection, the quantity of virus in this tissue is very low and does not influence the duration of fecal shedding [16, 20]. Several factors are involved in the degree and period of AIV shedding. Some studies have revealed that avian infectious bronchitis vaccines increased the duration and degree of AIV shedding and decreased the efficacy of H_9_N_2_ AIV vaccines in reducing AIV fecal shedding [21]. Some gross lesions, such as subcutaneous hyperemia and many petechial to ecchymotic hemorrhages, and stunting have been reported in chicken embryos inoculated with AIV+infectious bronchitis live vaccine virus [4].

Despite the decrease in the replication and shedding of the influenza virus, the clinical signs of the disease were not considerably different between the chickens of subgroups A and B. Costa-Hurtado et al. (2014) coinfected chickens and turkeys with lentogenic Newcastle disease virus and LPAIV and reported that the replication pattern of these viruses changed, -clinical signs were unaffected. The effects on viral replication varied according to the time of infection and species. The results suggested that infection with a heterologous virus may result in temporary competition for cell receptors or competent cells for replication and that this competition may be interferon-mediated and decreases with time [22]. Pantin-Jackwood et al. (2015) reported that co-infection with velogenic Newcastle disease virus and LPAIV in domestic ducks did not affect clinical signs but changed the pattern of virus shedding and transmission [13]. It appears that influenza symptoms are likely contributing to issues other than the degree of virus replication. However, suppression of AIV fecal shedding is the most interesting issue in this study. Chicken age, IBDV strain, and the time interval between inoculations of the two viruses may greatly affect AIV shedding. Vaccination against IBD at three weeks of age could be a practical approach to inhibit probable influenza infection.

## Conclusion

IBDV infection suppresses AIV shedding in older chickens. According to the results, infection with Gumboro virus at three weeks of age cannot be considered to exacerbate viral shedding or the symptoms of avian influenza.

## Acknowledgments

We thank Mr. Mahdi Asadsangabi, expert virology laboratory technician, and Mitra Mohammadi, master poultry science technician at the school of veterinary medicine, Shiraz University for their assistance that greatly improved the project.

## References

1. Sharma JM, Kim I-J, Rautenschlein S, Yeh H-Y. Infectious bursal disease virus of chickens: pathogenesis and immunosuppression. Developmental & Comparative Immunology. 2000;24(2-3):223–35.

2. Berg TPVD. Acute infectious bursal disease in poultry: a review. Avian pathology. 2000;29(3):175–94.

3. Montgomery RD, Maslin WR. Effect of infectious bursal disease virus vaccines on persistence and pathogenicity of modified live reovirus vaccines in chickens. Avian diseases. 1991:147–57.

4. Haghighat-Jahromi M, Asasi K, Nili H, Dadras H, Shooshtari A. Coinfection of avian influenza virus (H9N2 subtype) with infectious bronchitis live vaccine. Archives of virology. 2008;153(4):651–5.

5. Rosenberger J, Gelb Jr J. Response to several avian respiratory viruses as affected by infectious bursal disease virus. Avian diseases. 1978:95–105.

6. Fadly A, Winterfield R, Olander H. Role of the bursa of Fabricius in the pathogenicity of inclusion body hepatitis and infectious bursal disease viruses. Avian Diseases. 1976:467–77.

7. Yuasa N, Taniguchi T, Noguchi T, Yoshida I. Effect of infectious bursal disease virus infection on incidence of anemia by chicken anemia agent. Avian Diseases. 1980:202–9.

8. Giambrone JJ, Anderson WI, Reid WM, Eidson CS. Effect of infectious bursal disease on the severity of Eimeria tenella infections in broiler chicks. Poultry science. 1977;56(1):243–6.

9. Bano S, Naeem K, Malik S. Evaluation of pathogenic potential of avian influenza virus serotype H9N2 in chickens. Avian diseases. 2003;47(s3):817–22.

10. Alexander DJ. An overview of the epidemiology of avian influenza. Vaccine. 2007;25(30):5637–44.

11. Mosleh N, Dadras H, Mohammadi A. Molecular quantitation of H9N2 avian influenza virus in various organs of broiler chickens using TaqMan real time PCR. Journal of molecular and genetic medicine: an international journal of biomedical research. 2009;3(1):152.

12. Pan Q, Liu A, Zhang F, Ling Y, Ou C, Hou N, et al. Co-infection of broilers with Ornithobacterium rhinotracheale and H9N2 avian influenza virus. BMC veterinary research. 2012;8(1):104.

13. Pantin-Jackwood MJ, Costa-Hurtado M, Miller PJ, Afonso CL, Spackman E, Kapczynski DR, et al. Experimental co-infections of domestic ducks with a virulent Newcastle disease virus and low or highly pathogenic avian influenza viruses. Veterinary microbiology. 2015;177(1-2):7–17.

14. Wu C, Lin T, Zhang H, Davis V, Boyle J. Molecular detection of infectious bursal disease virus by polymerase chain reaction. Avian Diseases. 1992:221–6.

15. Falcone E, D’Amore E, Di Trani L, Sili A, Tollis M. Rapid diagnosis of avian infectious bronchitis virus by the polymerase chain reaction. Journal of virological methods. 1997;64(2):125–30.

16. Mosleh N, Dadras H, Mohammadi A. Evaluation of H9N2 avian influenza virus dissemination in various organs of experimentally infected broiler chickens using RT-PCR. Iranian Journal of Veterinary Research. 2009;10(1):12–20.

17. Karimi S, Dadras H, Mohammadi A. The effect of the extracts of Echinacea purpurea and Sambucus nigra (black elderberry) on virus shedding in H9N2 avian influenza infected chickens. Iranian Journal of Veterinary Research. 2014;15(3):256–61.

18. Meng S, Yang L, Xu C, Qin Z, Xu H, Wang Y, et al. Recombinant chicken interferon-α inhibits H9N2 avian influenza virus replication in vivo by oral administration. Journal of Interferon & Cytokine Research. 2011;31(7):533–8.

19. Raj GD, Rajanathan TC, Kumanan K, Elankumaran S. Changes in the cytokine and Toll-Like receptor gene expression following infection of indigenous and commercial chickens with infectious bursal disease virus. Indian Journal of Virology. 2011;22(2):146.

20. Kwon J-S, Lee H-J, Lee D-H, Lee Y-J, Mo I-P, Nahm S-S, et al. Immune responses and pathogenesis in immunocompromised chickens in response to infection with the H9N2 low pathogenic avian influenza virus. Virus research. 2008;133(2):187–94.

21. Tavakkoli H, Asasi K, Mohammadi A. Effectiveness of two H9N2 low pathogenic avian influenza conventional inactivated oil emulsion vaccines on H9N2 viral replication and shedding in broiler chickens. Iranian Journal of Veterinary Research. 2011;12(3):214–21.

22. Costa-Hurtado M, Afonso CL, Miller PJ, Spackman E, Kapczynski DR, Swayne DE, et al. Virus interference between H7N2 low pathogenic avian influenza virus and lentogenic Newcastle disease virus in experimental co-infections in chickens and turkeys. Veterinary research. 2014;45(1):1.

